# Predictive olfactory learning in *Drosophila*

**DOI:** 10.1101/2019.12.29.890533

**Authors:** Chang Zhao, Yves F. Widmer, Soeren Diegelmann, Mihai A. Petrovici, Simon G. Sprecher, Walter Senn

## Abstract

Olfactory learning and conditioning in the fruit fly is typically modelled by correlation-based associative synaptic plasticity. It was shown that the conditioning of an odor-evoked response by a shock depends on the connections from Kenyon cells (KC) to mushroom body output neurons (MBONs). Although on the behavioral level conditioning is recognized to be predictive, it remains unclear how MBONs form predictions of aversive or appetitive values (valences) of odors on the circuit level. We present behavioral experiments that are not well explained by associative plasticity between conditioned and unconditioned stimuli, and we suggest two alternative models for how predictions can be formed. In error-driven predictive plasticity, dopaminergic neurons (DANs) represent the error between the predictive odor value and the shock strength. In target-driven predictive plasticity, the DANs represent the target for the predictive MBON activity. Predictive plasticity in KC-to-MBON synapses can also explain trace-conditioning, the valence-dependent sign switch in plasticity, and the observed novelty-familiarity representation. The model offers a framework to dissect MBON circuits and interpret DAN activity during olfactory learning.

## Introduction

Predicting the future from sensory input is fundamental for survival. Co-appearing stimuli can be used for improving a prediction, or for predicting important events themselves, as observed in classical conditioning. In fruit fly odor conditioning, an odor that will become the conditioned stimulus (CS), is paired with the unconditioned stimulus (US), here an electroshock, that triggers an avoidance behavior and in internal representation of a negative value (valence). After conditioning, and the negative value representation – although not the full unconditioned response – previously elicited by the electroshock will be reproduced by the odor itself. Classical conditioning theories posit that throughout learning the odor becomes predictive for the electroshock^1–3^. During learning, the prediction error decreases, and learning stops when the predictive odor value matches the strength of the electroshock.

Predicitve olfactory learning in fruit flies is a widely recognised concept in the experimental literature^4, 5^, and dopaminergic neurons (DANs) in the mushroom body (MB) have been suggested to predict punishment or reward^6–8^. Yet, despite the acknowledgment of its predictive nature, computational models on fruit fly conditioning are mostly guided by the formation of associations, a notion that relates more to memories rather than predictions (for a recent outline of this controversy see^9^). Similarly, the concept of predictive learning is well recognized for olfactory conditioning in insects in general, but to our knowledge, synaptic plasticity models are not formulated in terms of explicit predictions, but rather in terms associations and correlations, with plasticity being driven by two or three factors, each representing a temporal nonlinear function of the pre- or postsynaptic activities or of a modulatory signal, sometimes combined with homeostatic plasticity. This type of associative models are exist for fruit flies ^10, 11^, locusts^12, 13^ or honey bees^14, 15^. They differ from target learning, where the unconditioned stimulus sets a target that is learned to be reproduced by the conditioned stimulus. Target learning becomes predictive learning when including a temporal component. It involves a difference operation, and learning stops when the target is reached. The stop-learning feature is difficult to be reproduced by purely correlation-based associative learning, while a purely predictive model intrinsically captures also associative properties.

Associative learning was suggested to be implemented through spike- or stimulus-timing dependent plasticity (STDP) that would underlay conditioning. STDP strengthens or weakens a synapse based on the temporal correlation between the US (electroshock) and the CS (odor), both on the neuronal time scale of 10’s of milliseconds^13^ and on the behavioral time scale of 10’s of seconds^7, 16, 17^. Whether an association is strengthened by just repeating the pairing until the behavioral saturation is reached^18^, or the association saturates due to a faithful prediction, however, has not been investigated in the fruit fly so far (Figure 1). Here we show that olfactory conditioning in *Drosophila* is better captured as predictive plasticity that stops when a US-imposed target is reached, rather than by correlation-based plasticity, such as STDP, that does not operate with an explicit error or a target. According to our scheme, it is only the aversive/appetitive value of the US that is predicted by the CS after faithful learning, not the US itself. Based on the common value representation in the mushroom body output neurons (MBONs, Figure 1), the corresponding avoidance/approach reaction as one aspect of the unconditioned response is elicited by the CS alone.

**Figure 1.**
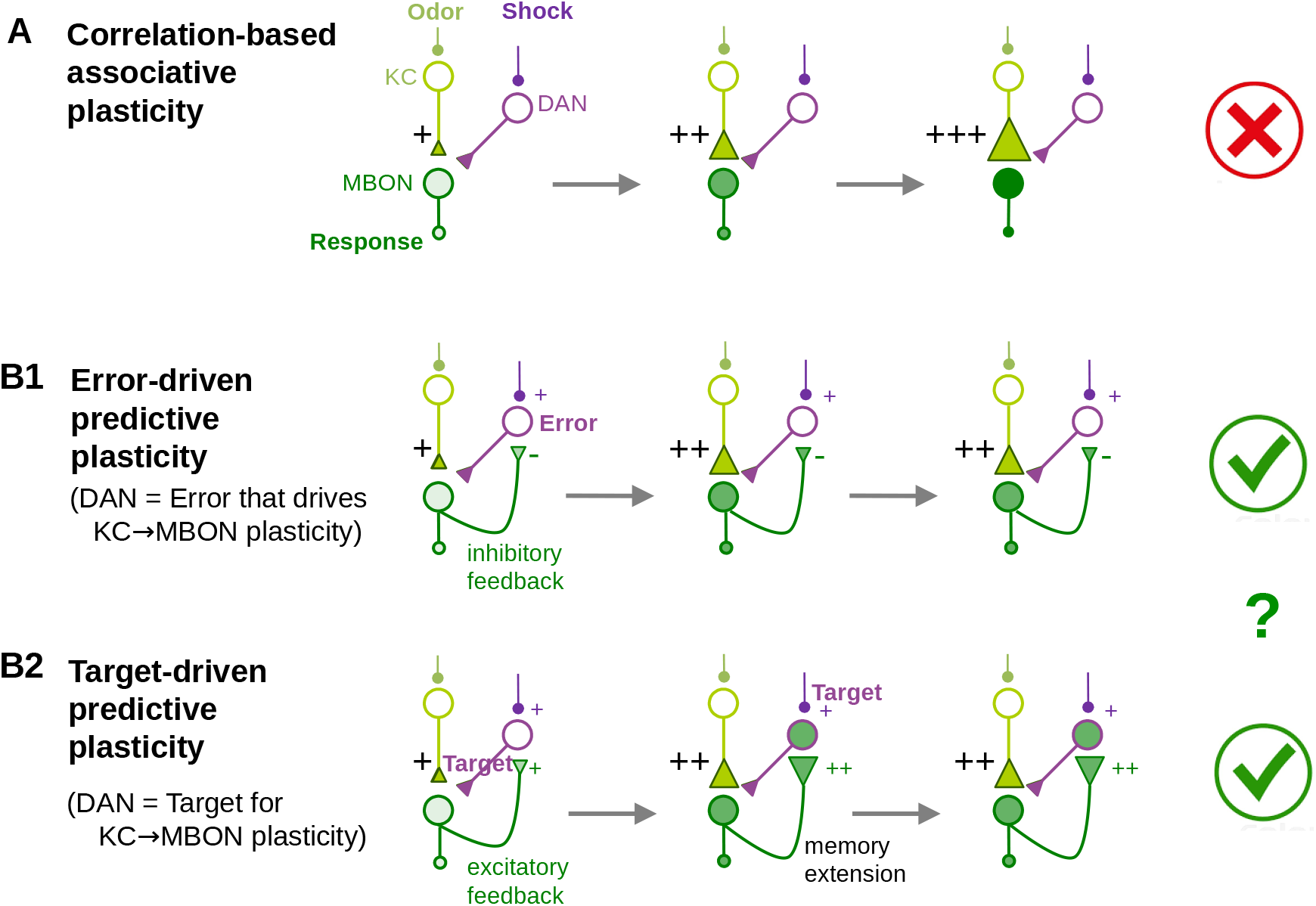
Associative versus error- or target-driven predictive plasticity. **(A)** Pairing of an odor (CS) with a shock (US) is typically thought to induce correlation-based synaptic strengthening of the synapses mediating the conditioned response, here from the Kenyon cells (KCs) to the mushroom body output neurons (MBONs). Repeated pairing always leads to a stronger association strength. **(B)** In predictive coding, plasticity stops when the strength of the US is correctly predicted by the CS. **(B1)** Plasticity of the KC-to-MBON synapses can be driven by the prediction error ‘US-CS’, formed by the dopaminergic neurons (DANs) that calculate the difference between the internal shock representation and the odor-evoked prediction by the MBONs. **(B2)** Synaptic plasticity can also be driven by a target for the MBON activity, set to the desired aversive value of the odor (i.e. the shock strength) and represented by the DAN activity. To extend the memory life time of the odor value, the DANs may themselves be driven by the recurrent MBON activity, and the MBON-to-DAN synapses may also be learned via target-driven plasticity to predict the shock strength. Our experiments and models exclude A and suggest further experiments to distinguish B1 and B2.

The *Drosophila* olfactory system represents a unique case for studying associative /predictive learning, and the MB is known to be essential in olfactory learning^18–21^. The Kenyon cells (KCs) receive olfactory input from olfactory projection neurons and form a sparse representation of an odor ^14, 22, 23^. The parallel axons of the Kenyon cells (KCs) project to the MB lobes^4, 24^, along which the compartmentalized dendritic arbors of the MB output neurons (MBONs) collect the input from a large number of KCs. Reward or punishment activates specific clusters of DANs PAM and PPL1, respectively which project to corresponding compartments of the MB lobes^25–27^, modulating the activity of the MBONs and the behavioral response^16, 28–30^. Recently, a detailed mapping of the MB connectome has been accomplished for larvae and of the vertical lobe for the adult Drosophila^31, 32^. Several studies show that not only the feedforward modulation from DANs to MBONs, but also the feedback from MBONs to DANs play an important role in olfactory learning^33, 34^.

Previous studies have given insights into the possible cellular and subcellular mechanisms of olfactory conditioning. Yet, the suggested learning rules ^10–14, 17, 35^ remain correlation-based and miss the explicit predictive element postulated by the classical conditioning theories^1, 2^. Here, we present distinctive conditioning experiments showing that olfactory learning is best explained by predictive plasticity (Figure 1). These experiments, in contrast, could not be reproduced by various types of correlation-based associative learning rules. A mathematical model captures the new and previous data on olfactory conditioning, including trace conditioning. The model encompasses the odor/shock encoding and the learning of the aversive odor value with the stochastic response. We further suggest how the predictive plasticity could be implemented in the MB circuit, with MBONs encoding the value (‘valence’) of the odor stimulus, and DANs calculating either the error or the target that drives the KC-to-MBON plasticity. The predictive plasticity rule for the KC-to-MBON synapses is shown to be consistent with the experimental results showing the involvement of these synapses in the novelty-familiarity representation.

## Results

### Model of the shock representation and the unconditioned response

Aversive odor conditioning is about learning to evoke the avoidance behavior by the conditioned odor alone, as it is evoked by the electroshock. Before describing the acquisition of the conditioned behavior we characterize the unconditioned behavior of the fruit flies.

#### Experiment

In the minimal shock detection experiments, fruit flies in a testing chamber had the choice between moving to either of two arms, with one arm being electrified (with voltage strength *S*) and the other not. After 30s we counted the number of flies in the electrified and non-electrified control arm, *N*_electr_ and *N*_nonel_, respectively (the few remaining in the testing chamber not being counted). The empirical performance index (PI) for the pure shock application without conditioning is defined as the relative difference, 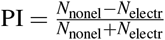. This empirical PI can be approximated by a theoretical PI that is a function of the stimulus strength *S*. When the stimulus strength is equal to the minimal strength *S*_∘_ that just elicits a behavioral response, the PI vanishes, PI(*S*_∘_) = 0, and for increasing stimulus strength the PI asymptotically tends towards 1. We parametrize

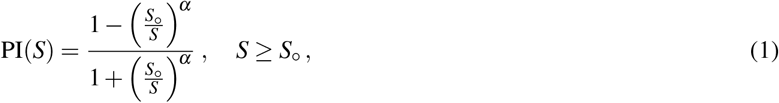

with sensitivity parameter *α* telling how steeply PI(*S*) grows from 0 at *S* = *S*_∘_ towards 1 for large *S*. We experimentally estimated the size of the minimal shock intensity to be *S*_∘_ ≈ 7*V* (Methods).

#### Shock representation

To explain the behavior as emerging from a neuronal representation we map the shock stimulus to hypothetical neuronal activities. Plasticity will then also be described in terms of these internal activities.

We assume an internal representation, *s*, of the electroshock following Weber-Fechner’s law^36^,

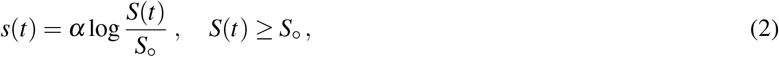

where *S*_∘_ and *α* are as introduced in Equation 1. For *S* < *S*_∘_ we set *s* = 0. Equation 2 yields a re-interpretation of the behavioral parameters *S*_∘_ and *α* that characterize the PI in terms of sensory ‘perception’: *S*_∘_ is the just detectable stimulus strength and *α* becomes the linear scaling of the sensory activity.

#### Unconditioned response

To describe the unconditioned response out of the internal representation, we consider the probability *p*_us_(*s*) of escaping from the shock stimulus (the US) given *s*. We first note that the PI can be expressed in the form PI = 2*p*_us_ − 1, with *p*_us_ = *N*_nonel_/(*N*_nonel_ + *N*_electr_) being the empirical frequency for an individual fruit fly to move to the non-electrified versus the electrified arm. With the avoidance probability of the form 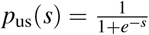, the PI becomes a function of the internal shock representation *s*, PI(*s*) = 2*p*_us_(*s*) − 1. This is consistent with the definition of PI(*S*) from Equation 1 as can be checked by substituting the expression for *s* given in Equation 2 (the performance index as a function of *s* is PI = (1 − *e*^−*s*^)/(1 + *e*^−*s*^)).

As an aside, one may also consider other mappings of the shock strength *S* to the internal representation *s* as this is not constrained enough by the data. For instance, one may argue that fruit flies perceive electric shocks following Steven’s power law, as it was originally measured in humans^37^. Steven’s law postulates that the internal representation would have the form *s* = (*S/S*_∘_)^*α*^ instead of the logarithmic Weber-Fechner law. From this representation the original behavioral response *p*_us_(*s*) is obtained when the readout from the state is of the form *p*_us_(*s*) = 1/(1 + *s*^−1^). The PI is then calculated according to PI(*S*) = PI(*s*) = 2*p*_us_(*s*) − 1 = (1 − *s*^−1^)/(1 + *s*^−1^).

### Odor conditioning depends on the temporal shock distribution

We next turn to the odor conditioning. It was previously investigated how the associative strength of the conditioned odor increases with the strength of the paired electroshock and the number of pairings, while saturating at some level, with respect to both the shock strength and the paring repetition^18^. We asked whether these saturation effects originate from behavioral limitations, or whether they originate from a quick and faithful learning of the intrinsic value of the electroshock strengths.

To address this question, we differently packaged the total of 100V into 1*100V, 2*50V, 4*25V and 8*12.5V electroshocks and asked whether the repeated smaller shock strengths (8*12.5V) would lead to premature saturation that would then not be caused by behavioural limitations but rather by some dedicated learning behavior. We distributed these shock packages across the 60s odor presentation time (Figure 2A, Methods), and let the fruit flies choose during 120s between a conditioned and neutral odor. The learning index (LI) that characterizes the conditioned response is defined analogously to the PI by the relative number of fruit flies that choose the unconditioned control odor (*N*_CS−_, more precisely, the odor that was conditioned with zero shock strength) versus the conditioned odor (*N*_CS+_), i.e.

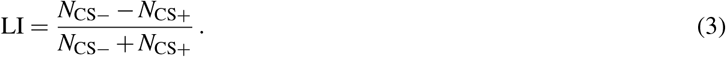

**Figure 2.**
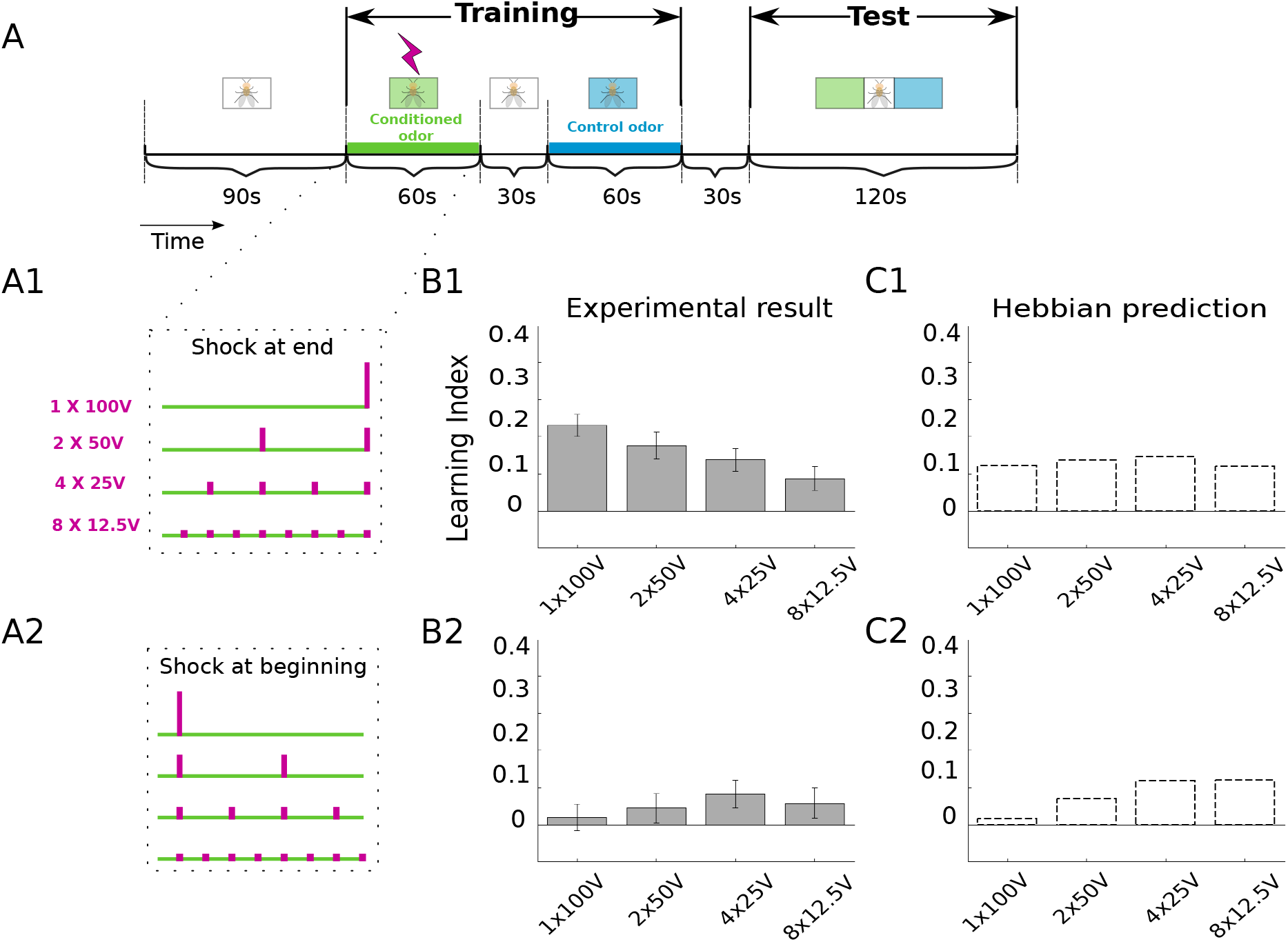
Temporal sequence conditioning is not fit by Hebbian plasticity. **(A)** Experimental protocol as explained in the main text, with total shock strength of 100 V distributed in time across the 60 s exposure to the conditioning odor (alignment towards the end, A1, and towards the beginning, A2). The duration of an individual voltage shock (a magenta vertical bar) is 1.5 s. **(B)** Experimental results showing the LI for the corresponding shock distributions in A1 and A2, respectively. For shocks at the end, the LI decreases with decreasing individual shock strength. Error bars represent the standard error of the mean (SEM). **(C)** Fit by the Hebbian plasticity, Equation 6. For shocks at the end, a roughly constant LI is produced. For the optimized parameters we extracted *S* = 7 V from the minimal shock detection experiment (Figure 2-1), *τ_o_* = 15 s from trace conditioning experiments^28^, and optimized the product *αη* = 0.0723.

The LIs gradually decreased with decreasing electroshock strength if the shocks were applied towards the end of the odor presentation time, and the additional repetitions of the weaker electroshocks could not revert this trend (Figure 2B1). Yet, when the same shocks were distributed towards the beginning of the odor presentation time, the LIs remained small, with a tendency to increase with decreasing electroshock strength (Figure 2B2).

The avoidance behavior depends in a complicated way on the shock strengths and the shock timings. To explain these behaviors we next formalize the value representation, the decision making, and two different types of plasticity models.

### From internal value representation to stochastic responses

The basic observation of conditioning is that, after long enough conditioning time, the conditioned behavior eventually mimics the unconditioned behavior. In our model this implies that the learning index LI converges to the performance index PI (Equation 1). We assume that at any moment in time, a presented odor elicits some activity *o* in the KCs that reflects the odor intensity. In the experiments, an odor was either present or absent, and hence *o*(*t*) = 1 or 0. The considered MBON activities are assumed to represent the aversive value (*v*) of the odor. As MBONs are driven by KC, we postulate that the MBON activity takes the form

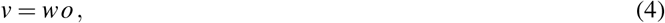

where *w* is the synaptic strength (weight) from the KCs to the MBONs.

#### Model of stochastic action selection

The conditioned response upon odor stimulus appears to be stochastic for an individual fruit fly. It is therefore modeled in terms of the avoidance probability that itself depends on the MBON activity. For simplicity we postulate that this avoidance probability in response to the conditioned stimulus (CS, the odor), has the same form as the one to the unconditioned stimulus introduced above, *p*_us_(*s*), but with *s* replaced by *v*,

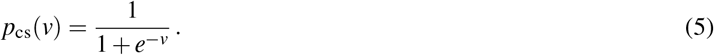

As it is for the PI, the LI can be expressed in terms of this avoidance probability, LI(*v*) = 2*p*_cs_(*v*) − 1. Remember that fruit flies may remain in the test tube (estimated to be less than 5%) and that the LI is calculated based on the fruit flies that effectively moved to one of the two chambers (Equation 3). Hence, the interpretation of *p*_cs_(*v*) on the level of the individual fly is, strictly speaking, the conditional probability that, given the fly ‘decides’ to move, it actually moves away from the conditioned odor.

The model postulates that the decision for each individual fruit fly is a stochastic (Bernoulli) process that only depends on the current MBON activity *v* = *wo*, and in particular does not depend on previous decisions. In fact, when re-testing the population of fruit flies that escaped from the odor in a first test trial (a fraction *p*_cs_ of the overall test population), the same fraction *p*_cs_ of this sub-population escaped again in a second test trial, despite the putative extinction of memory caused by the first test (see the Test-Retest experiment in^38^). Intriguingly, when waiting 24h so that the first conditioning was forgotten, conditioning the successfully escaped and the unsuccessfully non-escaped flies from the first conditioning experiment separately again, the same LI was achieved by both groups. This shows that not only a single response is stochastic, but also the learning (see again^38^, cross-checked by us for a 8 × 12.5*V* stimulation, results not shown). A statistical evaluation of the model with the same number of flies (*N*_fly_) and trials (*N*_trial_) as in the experiment gives equal or smaller variance in the LI of the model fruit flies as compared to the experiment (Figure 3A1). This implicitly quantifies additional sources of stochasticity in the experimental setup or in the individual fruit fly that have already been described in honeybees^39, 40^ and that go beyond our 1-state stochastic Markov model.

**Figure 3.**
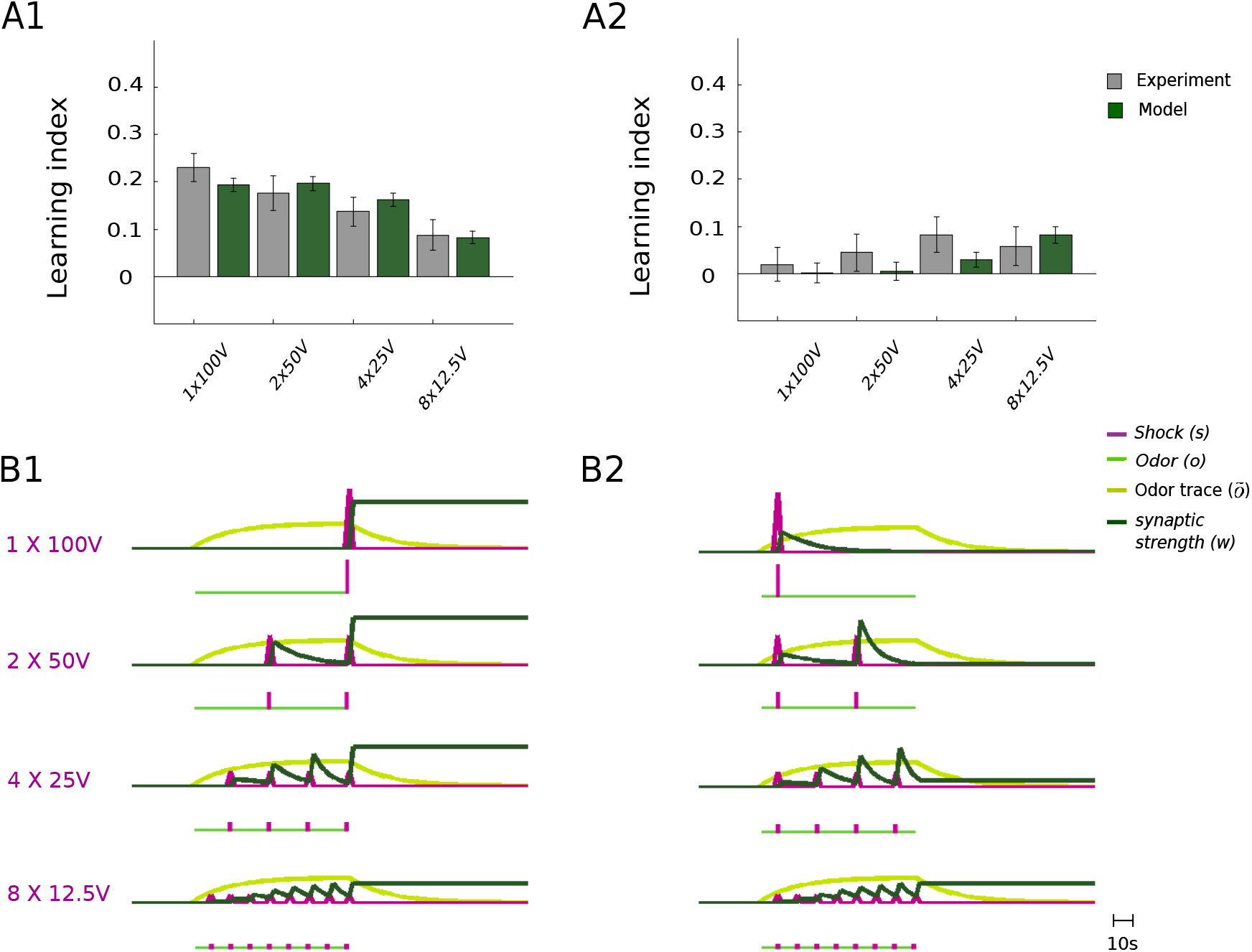
Predictive plasticity captures the sequence conditioning experiments. **(A)** In contrast to the purely associative learning (Equation 6), the predictive plasticity (Equation 7) fits well the shock-at-end data (A1) and the shock-at-beginning data (A2, see Figure 2). Error-bars represent SEM for both data and model. In the model, stochasticity enters through the Bernoulli process according to which each of the model flies chooses to avoid (with probability *p*_cs_(*v*), Equation 5) or approach (with probability 1 *p*_cs_(*v*)) the odor. The same number of flies, *N*_fly_, and the same number of trials, *N*_trial_, was used in the model as in the experiment. **(B)** Traces for odor (*o*), shock (*s*), and synaptic strength (*w*) for the odor-to-shock prediction during conditioning, for the shock-at-end (B1) and shock-at-beginning (B2) protocols. Note that between the shocks, *w* decays towards 0 as the target for *v* = *wo* is *s* = 0 according to the predictive learning rule. The weight does not change if the prediction matches the shock representation, *wo* = *s*, e.g. when both are 0. The optimized parameters are: *S*_∘_ = 6.90 V, *α* = 0.79, *τ_o_* = 14.25 s, Δ*η* = 0.057, *τ_η_* = 133.48 s, with mean square error MSE = 6.393 × 10^−4^ across all experiments (including the ones below).

The model of stochastic action selection expressed by Eq. 5 assumes that there is only one stimulus type present, either the odor (CS) or the shock (US), and the odor triggers the avoidance reaction with probability *p*_cs_(*v*), and the shock with probability *p*_us_(*s*). The experiment may also be setup such that in one arm of the test chamber the CS and in the other the US is present, and the fruit fly can decide whether to move at all or not, for instance, as studied in^41^. In this case the probability of moving in neither of the two arms depends on the difference between the CS- and US-induced value, 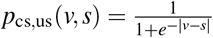, and this probability may be represented downstream of the MB, as also suggested in^41^. Alternatively, the DAN may represent the US and the MBON may depend on both the CS and US, along the lines of the wiring scheme for the target-driven predictive plasticity outlined below (Discussion with Figs 1B & 6B).

### Associative learning models do not fit the conditioning data

Learning is suggested to arise from appropriately modifying the strength *w* of the KC-to-MBON synapses. The synaptic modification affects the aversive value of the odor following the linear relation *v* = *wo* (Equation 4), and this determines the conditioned response given by the escape probability *p*_cs_(*v*), see Equation 5.

The common conception of conditioning is that the associative strength, *w*, is changed proportionally to some nonlinear functions of the pre- and postsynaptic activities, possibly modulated by a third factor. To exemplify the essence of associative learning, although this may not do justice to the more complex cited models, we consider a simplified version where the synaptic weight change is proportional to both the strengths of the unconditioned and the conditioned response,

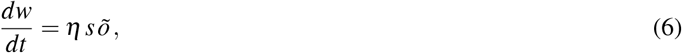

with proportionality factor *η* defining the learning rate (cf. Figure 1A). Here, *õ* is the low-pass filtered odor *o* that follows the dynamics 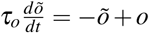, with a time constant *τ_o_* being on the order of ten seconds. It can be interpreted as a presynaptic eligibility trace that keeps the memory of the presynaptic activity, here the odor *o*, to be associated with the postsynaptic quantity, here the shock representation *s*.

This simple Hebbian rule (Equation 6) is not able to fit the sequential conditioning data. In fact, for the shocks at the end, the Hebbian model roughly shows the same LI for the weak and strong stimuli, as it were the total stimulus strength that would count (Figure 2C1). The concavity of the logarithmic shock representation by itself would rather favor an increasing LI for the repeated weaker stimuli 8*12.5V as compared to the 1*100V.

We considered a perhaps oversimplified Hebbian learning rule, 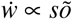, as one example of associative plasticity. To consider more sophisticated associative learning rules, we define synaptic weight changes that are functions of the correlation between odor- and shock-induced activity. We also tested these more general forms of associative learning that are based on linear and nonlinear functions of CS-US correlations, such as stimulus-timing dependent synaptic plasticity (STDP) of the form 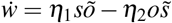, with 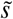 being the low-pass filtered *s* and *η_i_* arbitrary scaling factors. STDP, even after introducing nonlinearities, and also the covariance rule of the form 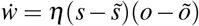, did all give roughly a 10 times worse fit (in terms of the MSE, Methods and Figure 3-1) than predictive plasticity explained next.

### Model of predictive plasticity

The failure of associative learning rules in reproducing our conditioning experiments can be corrected by adding an anti-Hebbian term of the form −*võ* to the rule 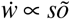, leading to

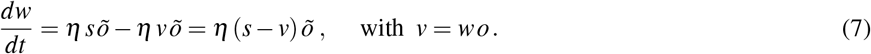

We interpret this combined Hebbian / anti-Hebbian plasticity rule as error correcting, with the difference between the shock and odor representation, *s* − *v*, as internal error.

This error-correction learning rule has a long history in the theory of neural networks where it first appeared as Widrow-Hoff rule^42^ that was extended to a temporal difference rule^43^, and recently reinterpreted in terms of dendritic prediction of somatic firing^44, 45^. It also relates to the predictive rule of Rescorla-Wagner^1^ previously applied to explain various fruit fly conditioning experiments^46^, although without considering a time-continuous learning scenario and the related temporal aspects. According to this rule, learning stops when the aversive value of the odor, *v*, predicts the internal representation of the shock stimulus, *v* = *s*. During predictive learning, when the synaptic eligibility trace is active, *õ*(*t*) > 0, the synaptic strength *w* is adapted such that the odor value converges to the internal shock representation, *v*(*t*) = *wo*(*t*) −→ *s*(*t*), with *s*(*t*) > 0 when the electroshock-voltage is turned on and *s*(*t*)=0 else. Correspondingly, the conditioned response converges to the unconditioned response, *p*_cs_(*v*) −→ *p*_us_(*s*). Crucially, during the time when the US is absent, *s* = 0 (while *õ*> 0), a neutral response is learned. On a behavioral level this appears as forgetting the shock prediction, and it also relates to the phenomenon of extinction in classical conditioning^1^.

To fit the conditioning experiments with ongoing electroshock-voltage we need to consider a learning rate that adapts in time. Learning speeds up when the strength of the voltage increases. A stronger voltage triggers initially a higher learning rate that, with ongoing voltage stimulation, decays with a time constant *τ_η_* on the order of 2 min. A stepwise increase of *s* by Δ*s* (as it appears at the onset of an electric shock) leads to a stepwise increase of the initial learning rate *η* by Δ*η* Δ*s* for an optimized parameter Δ*η* (Methods).

In contrast to the pure associative rule, the predictive rule (Equation 7) qualitatively and quantitatively reproduces the conditioning experiments (Figure 3). With the predictive learning rule, the 1*100V pairing at the end of the odor presentation elicits the strongest conditioned response, while the response is much weaker after the distributed 8*12.5V pairing, as also observed in the experiment. The reason is that the synaptic weight *w* decreases between the shocks while the odor is still present (green traces in Figure 3). As in the extinction experiments, the presence of the CS alone leads to the prediction that no US is present, and hence to an unlearning of the previously acquired US prediction.

### Repetitive and ongoing conditioning reveals its predictive nature

To bolster our hypothesis that olfactory conditioning in the fruit fly is predictive rather than associative, we further tested the model to repetitive and continuously ongoing odor-shock pairings. If the hypothesis is correct, during repeated or extended pairing, learning should in both cases stop when shock strength is correctly predicted by the odor. In particular, the learning performance is expected to saturate at a level below the maximally achievable performance. This is in fact what we observed.

When repeating the previously described block of 4*25V conditioning shocks with 15s inter-shock-intervals, the LI showed a saturation after a single block (Figure 4A,B). When conditioning with half of that block, i.e. with only 2*25V conditioning shocks in 15s, roughly 70% of the saturation level is reached. The same repetition experiment was performed with 4*50V pulses, confirming that also for a stronger US the LI quickly saturated (Figure 4C). Again, neither the pure associative rule, nor the covariance rule or the more sophisticated STDP rules, could reproduce this data (Figure 3-1).

**Figure 4.**
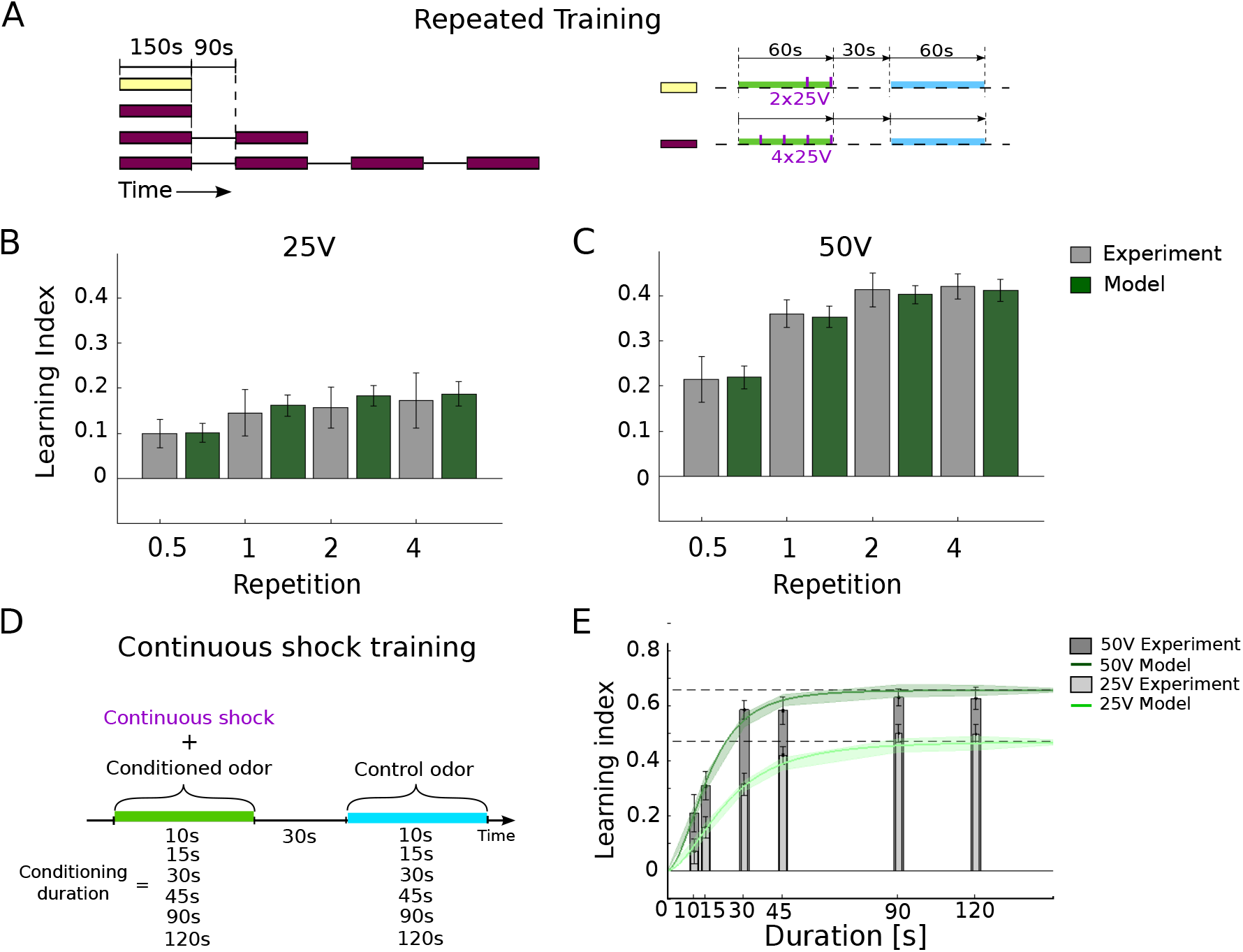
Repetitive and ongoing conditioning is captured by the predictive plasticity. **(A)** The repeated training consisted of 1, 2, 4 repetitions of a standard training block (dark red bar, as in Figure 3, A&A1) composed of 4*25V shocks, followed by a break and a control period. Half of a training block was considered with 2*25V shocks towards the end of the 60s odor presentation (yellow bar). **(B)** The LI saturates after a full block (1 Repetition, gray), as also reproduced by the predictive plasticity model (green). **(C)** The same protocol with the same number of shocks as in (B), but with 50V instead of 25V shocks. A second training repetition did only slightly increase the LI and for further repetitions it again remains constant. This is reproduced by the predictive plasticity, but not by the various associative plasticity models (see Supplementary Materials). **(D)** Protocol of ongoing odor-shock pairing, with voltages turned on during the full odor presentation time of 10s, 15s, 30s, 45s, 90s and 120s, both for 25V and 50V. **(E)** The LI for the time-continuous pairing saturates with a time constant of roughly 20s for the 50V and 30s for the 25V odor-voltage pairings. Predictive plasticity captures this saturation, with LI(*v*) converging towards LI(*s*) (dashed lines, Equation 8), for both the 25V and the 50V pairings.

An even more challenging test for the predictive learning rule is an odor-shock pairing where the electric voltage (either 25V or 50V) is turned on throughout the odor presentation time, from 10s up to 120s. After roughly 1 min of ongoing pairing the LI saturated, both in the data and the model (Figure 4D). In the model, learning saturates when the value *v* of the odor correctly predicts the shock, *v* = *s*, as expressed by a successful predictive learning (i.e. when learning ceases, 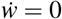, see Equation 7). During learning, when the value of the odor converges to the shock representation, *v* → *s*, the LI converges to the PI (as defined in Eqs 3 and 1),

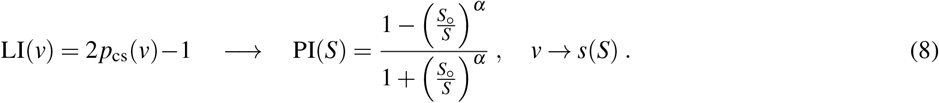

The equation is obtained from substituting *v* by *s* in the expression for *p*_cs_(*v*), Equation 5, and making use of Weber-Fechner’s law translating the shock strength *S* into the internal representation *s* (Equation 2). For our simple predictive plasticity model the exposure time to acquire the final performance can be explicitly calculated, and it is shorter for stronger ongoing voltage stimuli (Figure 4E and SI, Fig. 4-1).

### Trace conditioning is also predictive

Odor conditioning has also been studied in the form of trace conditioning (e.g.^28, 47^). A further test of our model is to apply it to these experiments, with the same parameters found to fit our data from Figure 3 and Figure 4. In trace conditioning, the electroshock is applied with a variable inter-stimulus-interval (ISI) after the onset of the odor presentation, and this ISI can even extend beyond the presentation time of the odor (Figure 5A). We considered the experimental protocol with 10 s odor presentation and an ISI varying from 5 to 30s, after which 4 conditioning electroshocks of 90V were applied with 0.2 Hz^28^. The LI gradually decreased with the length of the ISI, with a decay time of roughly 15 s. The model captures this phenomenon because the odor trace, entering as synaptic eligibility trace (*õ*) in the predictive plasticity rule, is still active for a while after the odor has been cleared up (Figure 5B). The identical set of 5 parameters has been used that were extracted from the previous experiments (*S*_∘_, *α*, *τ_o_*, Δ*η*, *τ_η_*, see caption of Figure 3).

**Figure 5.**
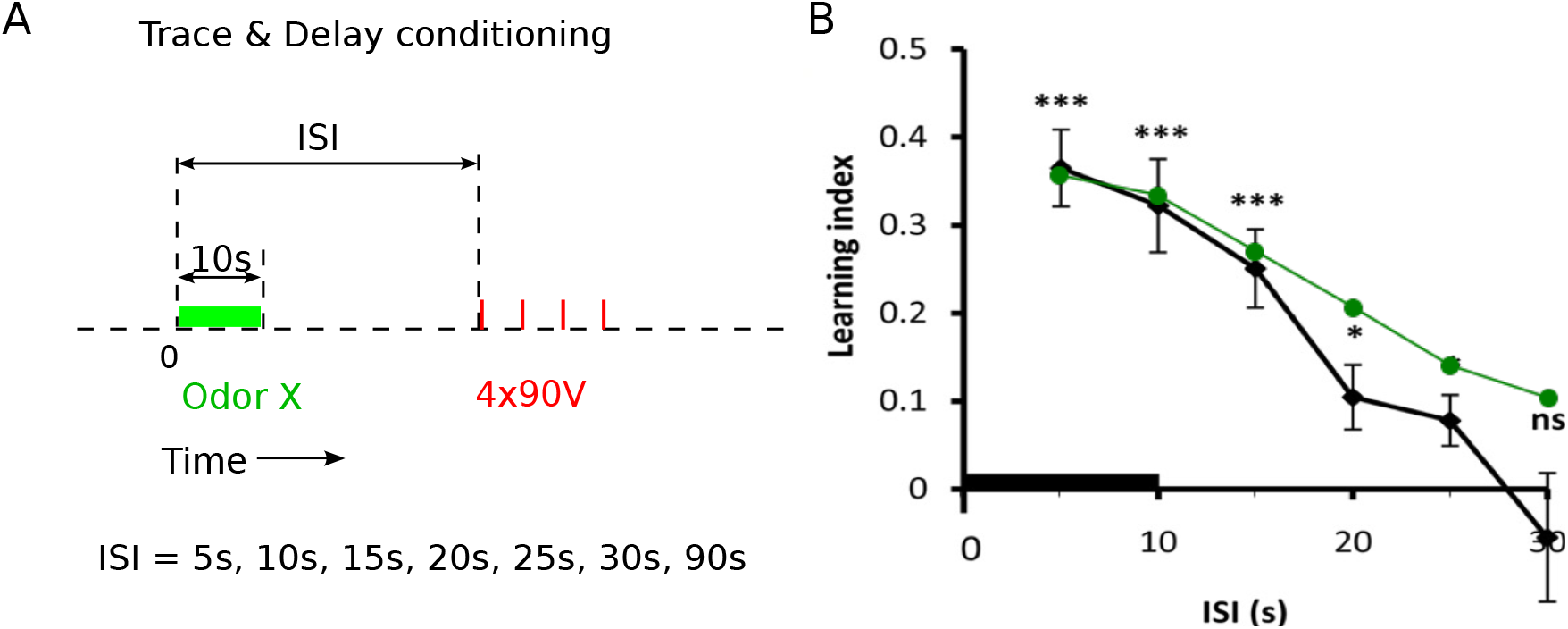
Trace conditioning is faithfully reproduced. **(A)** Experimental protocol of trace conditioning, with variable Inter-Stimulus Intervals (ISIs) from the onset of the odor (green) to the onset of the electroshock train (4 90V, 0.2Hz, each 1.25s, red bars). **(B)** The LI tested immediately after the conditioning with the different ISIs (reproduced from^28^). The model roughly captures this data (green line) without additional fitting of the parameters.

### MB circuits for error- or target-driven predictive plasticity

Based on anatomical connectivity patterns and previous plasticity studies we suggest two forms of how the predictive learning may be implemented in the recurrent MB circuit, via error- and target-driven predictive plasticity (Figure 1B). In both versions, learning is mainly a consequence of modifying the KC-to-MBON synaptic boutons^4, 30, 33, 48, 49^, but the role of the DANs is different. While the KC-to-MBON connections drive the MBONs based on the odor representation in the KCs, the shock information is provided by the DANs and gates the KC-to-MBON plasticity (see also^4, 7, 24, 25, 27^). The DANs themselves may either represent the error or the target for KC-to-MBON plasticity.

In the first implementation (error-driven predictive plasticity), the DANs themselves represent the prediction error *e* = *s − v*. They may extract this error from the excitatory shock input, *s*, and the inhibitory MBONs feedback providing the aversive value *v* of the odor (^33^, see Figure 6A1). The modeling captured the effect of learning on the behavioral time scale. To predict specific activity traces in the MB on a fine-grained temporal resolution we introduce the dynamics of the MB neurons. In the case of the DANs as error representation, the firing rates of the MBONs (*v*) and DANs (*e*) is given by

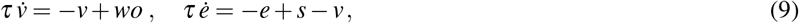

with a neuronal integration time constant *τ* in the order of 10 ms (Figure 6B1). The plasticity of the synapses from the KCs to the MBON is then driven by the DAN-represented prediction error *e* at any moment in time, 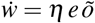, consistent with the predictive plasticity rule (Equation 7). Note that in the steady state, the DAN activity exactly represents the difference between the shock strength and its odor-induced prediction, *e* = *s* − *v*. After successful learning, the MBONs accurately match the shock representation and the DAN activity vanishes, *v* = *s* and *e* = 0.

**Figure 6.**
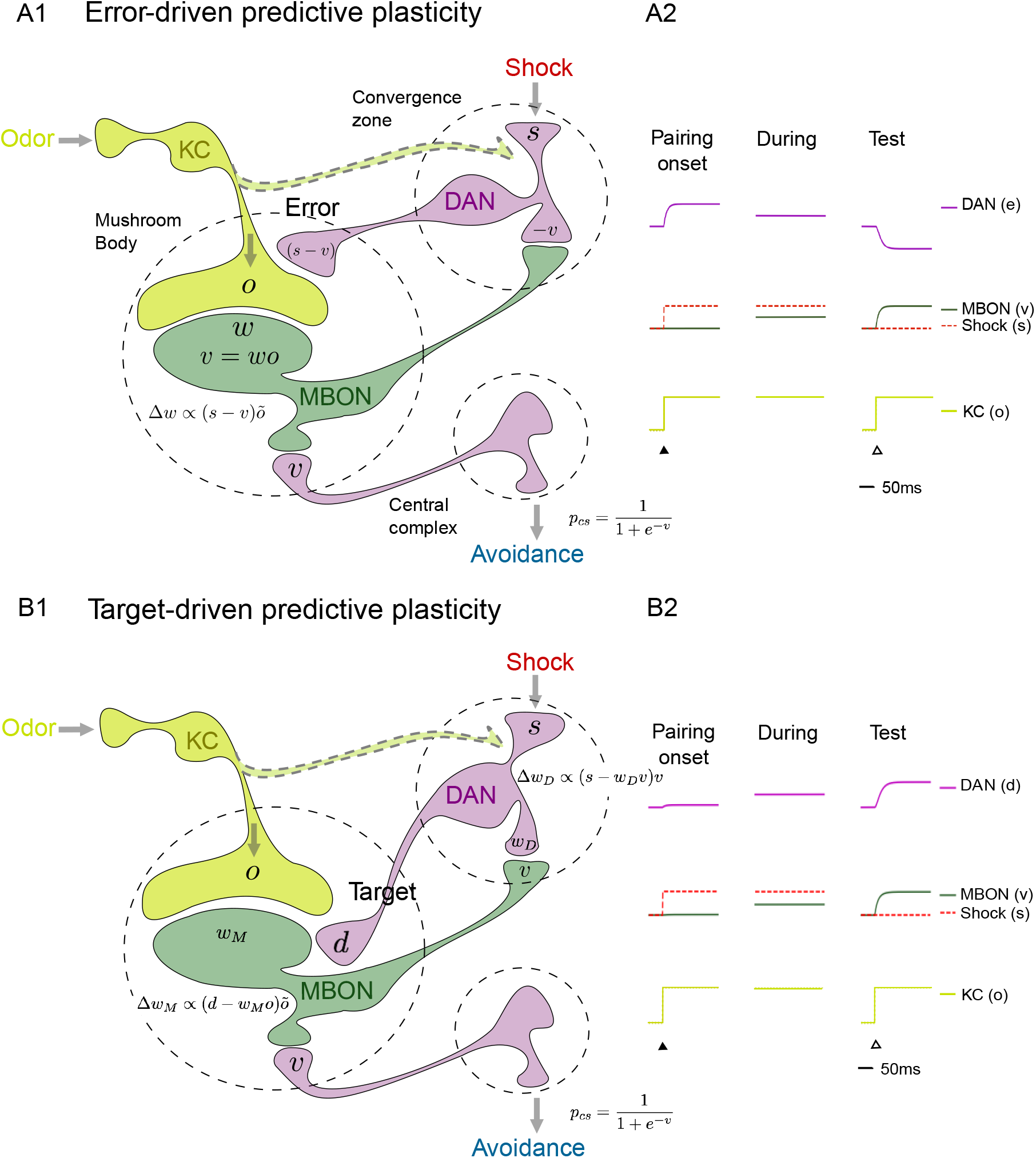
Suggested implementation of error- and target-driven predictive plasticity. **(A1)** Mushroom body circuits for olfactory error-driven predictive plasticity. Kenyon cells (KCs) carrying the odor information project to mushroom body output neurons (MBONs) through synapses encoding the aversive value (*v*) of the odor. The input triggered by the electroshock, *s*, drives the dopaminergic neurons (DANs) that are also inhibited by the MBONs. The DANs represent the prediction error, *e* = *s* − *v*, and modulate the KC-to-MBON synapses according to 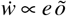, with *õ* representing the odor eligibility trace. The conditioned response probability (*p*_cs_, avoidance reaction) is a function of *v*. **(A2)** Neuronal activity traces of a DAN (*e*, magenta), a KC (*o*, light green) and a MBON (*v*, dark green), shown at the onset of an odor-shock pairing (‘Pairing onset’, full triangle), 20 s later (During), and later at the test when only the odor is presented (‘Test’, open triangle, cf. Eq. 9). The aversive value *v* steadily increases (dark green), while the prediction error, *e*, decreases throughout learning and becomes negative when the odor is presented alone (purple). **(B1)** Mushroom body circuit for target-driven predictive plasticity. Beside the shock stimulus, the DANs can also indirectly be excited by the MBONs (or directly by the KC, not shown) to form a shock prediction also in the DANs and prevent fast extinction. The shock stimulus (*s*) sets the target for the MBON-to-DAN plasticity, and the DANs (*d*) set the target for the KC-to-MBON plasticity (cf. Eq. 11). **(B2)** As in A2, but since the DANs now form a prediction of the shock itself based on *v*, their activity increases throughout learning, and they are also activated during the Test, when the conditioned odor is presented alone (Eq. 10). Sketch adapted from^33^ and^6^, where also excitatory feedback to the DANs is favored, as in version B.

In the alternative implementation (target-driven predictive plasticity), the DANs provide the learning target to the KC-to-MBON synapses while themselves being driven by the MBONs (Figure 6A2). These MBON-to-DAN synapses are also plastic and learn to predict the shock stimulus, just as the KC-to-MBON synapses do. A benefit of this recurrent prediction scheme is that the memory life time of the odor-shock prediction is extended. If after successful learning the odor is presented alone, the target for the KC-to-MBON plasticity is still kept at the original level via MBON-to-DAN feedback, and extinction of the shock memory slows down. The recurrent circuitry between MBONs (*v*) and DANs (with activity *d* instead of *e* to indicate that the DANs no longer represent the error but the target for the MBON learning) now becomes

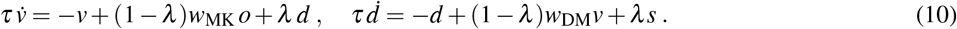

Here, *w*_MK_ and *w*_DM_ are the synaptic weights of KC-to-MBON and MBON-to-DAN synapses, respectively, and *λ* = 0.1 is the nudging strength of the postsynaptic teaching signal^44^). Both KC-to-MBON and MBON-to-DAN synapses follow the same form of error-correcting plasticity as in Equation 7,

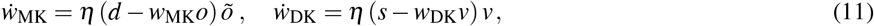

where the DAN activity *d* now serves as the target for KC-to-MBON synapses, while the shock stimulus *s* is the target for MBON-to-DAN synapses.

After successful learning, the activity of MBONs and DANs both predict the shock stimulus, *v* = *d* = *s* (as derived from the steady states of Eq. 11, see also Figure 6B2). If the shock stimulus is absent (*s*=0) during the presentation of the conditioned odor *o*, and the odor was previously conditioned to a shock strength *s*_∘_ while the DAN activity was fully learned (implying *w_D_* = 1), the MBON activity, supported by the recurrent DAN activity, becomes 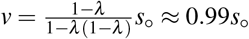 (as derived from the steady states of Eq. 10 with *λ* = 0.1, see Figure 6B2, column ‘Test’). Hence, the value of the odor faithfully predicts the conditioned shock strength also in this target-driven learning circuitry. Note that in the target-driven plasticity the KC-to-MBON plasticity 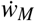 does not directly relay on the MBON activity since the activity target is imposed by the DAN’s, not by the MBON’s (Eq. 11).

#### Outlook: valence learning and novelty-familiarity representation

The concept of predictive learning can be extended to valence learning where each MBON represents a positive or negative valence, *v*^±^, coding for an appetitive and aversive value of a stimulus, respectively^24, 30, 50, 51^. For each valence, a specific cluster of DANs is involved in the sensory representation, PAM for positive and PPL1 for negative valences^4, 52^. In the full MB circuit the DANs further receive excitatory drive from the KCs (^32^, dashed connection in Fig. 6, here abbreviated by *w*_DK_), and the feedback circuit modulates the plasticity of the KC-to-MBON synapses^48^. The activities of the two valence classes of DANs can be modeled as in Eq. 10, but with multimodal input from the unconditioned appetitive or aversive stimuli (*s*^±^) and the odor representation in the KCs (*w*_DK_*o*). Together with the feedback from the corresponding MBONs via weights *w*_DM_, and introducing a saturating nonlinear transfer function *ϕ*, the DAN activities for the two valence clusters become

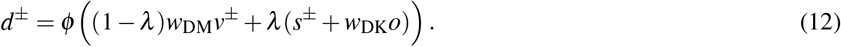

Plasticity in MBONs is known to be sign flipped when changing the valence of the stimulus^7, 30, 52^. This can be captured in the predictive plasticity model by imposing 0 as target when the stimulus and MBON valence do not match. For positive valence MBONs, the target can be set to *d* = *d*^+^(1 − *d*^−^), assuming that the DAN activities are restricted to the range between 0 and 1; for negative valence MBONs the target is *d*^−^(1 − *d*^+^). When a previously appetitively conditioned odor is now presented (*w*_MK_*o* > 0 for a positive valence MBON), together with a shock (*d* = 0), the postsynaptic error term in the learning rule now becomes negative, (*d* − *w*_MK_*o*) < 0, and the synapses get depressed rather than potentiated as in the first conditioning (Eq. 11).

The sign of the KC-to-MBON plasticity can also be changed in other ways. It has been shown that the familiarization to odors can depress MBON responses (in the *α*′3 compartment), while the response to previously familiarized stimuli is recovered^49^. To explain this phenomenon we extend the predictive plasticity to involve a partial redistribution of the total synaptic strength across the KC-to-MBON synapses, formally expressed by

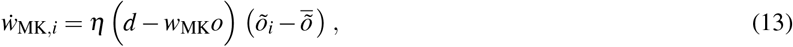

where we introduced a down-shift in the presynaptic term by the mean odor that exceeds the spontaneous activity level, 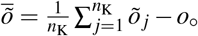. Here, the average is across all *n*_K_ Kenyon cell synapses, and we assume a spontaneous but sparse KC activity *o*_∘_ such that in average the activity of KC *i* satisfies *o_i_* ≥ *o*_∘_^53^. The spontaneous KC maps to the eligibility trace that is strictly positive, *õ_i_* ≥ *o*_∘_, and some spontaneous DAN activity *d*_∘_ inherited from the KCs, such that *d* ≥ *d*_∘_. Because (PPL1−*α*′3) DAN activity is necessary to observe repetition suppression^49^, we postulate that the learning rate is modulated by the DAN activity, *η* = *dη*_∘_, for some base learning rate *η*_∘_.

The various plasticity features of the KC-to-MBON synapses investigated in^49^ are consistent with the extended learning rule in Eq. 13. Repeated odor-evoked KC activation causes synaptic depression, assuming that odors are dominantly activating KCs and MBONs, but less so DANs, *d* < *w*_MK_*o* (here enters the saturation of the nonlinearity *ϕ* in Eq. 12), leading to the observed repetition suppression 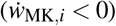 and explaining the behavioral familiarization of the flies to odors. The repetition suppression may depress the KC-to-MBON synapses such that in response to spontaneous KC activity (*o*_∘_) the MBON activity is now smaller than the spontaneous DAN activity, *w*_MK_*o*_∘_ *< d*_∘_. In the absence of an odor, the depressed KC-to-MBON synapses will therefore recover due to the spontaneous KC activity (Eq. 13), such that eventually the equilibrium is reached again when the spontaneously induces MBON activity matches the spontaneous DAN activity, *w*_MK_*o*_∘_ = *d*_∘_. This explains the ‘passive’ recovery of the MBON responses after odor familiarization^49^.

Further experimental investigations of the KC-to-MBON plasticity shows that optogenetically activating DANs alone potentiates the synapses. In our model this DAN-induced potentiation arises since for the isolated optogenetic DAN activation we have to assume that *d* > *w*_MK_*o*, and the presynaptic term in the plasticity rule (Eq. 13) is positive in average due to the spontaneous KC activity, 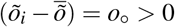. Next, if we assume that the optogenetic co-activation of MBONs (*v* > 0) and DANs (*d* > 0) applied in^49^ is such that *d < v* = *w*_MK_*o* (but with an increased learning rate *η* = *dη*_∘_), then the KC-to-MBON synapses get depressed, as reported from the experiment.

Finally, due to the partial weight redistribution, the repetition suppression during the familiarisation to a new odor implies the potentiation of the other synapses that are not activated, among them most of the previously suppressed synapses that were involved in the representation of a preceding odor. The reason for this heterosynaptic potentiation in our model is that repetition suppression is caused by a negative postsynaptic factor (*d* − *w*_MK_*o*) < 0 in Eq. 13 as explained above, implying the depression of an active synapses *i* for which 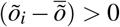, but also implying the potentiation of not activated synapses, since for those 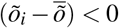 and hence 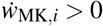. This odor-induced potentiation in other synapses explains the ‘active’ recovery from the repetition suppression as seen in the experiment^49^. Technically, 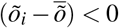 holds for not activated synapses only if we assume that the odor-evoked average activity in the KCs is well above the spontaneous activity level such that 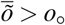.

## Discussion

### Predictive, but not correlation-based plasticity, reproduces experimental data

We reconsidered classical odor conditioning in the fruit fly and presented experimental and modeling evidence showing that olfactory learning, also on the synaptic level, is better described as predictive rather than associative. The key observation is that repetitive and time-continuous odor-shock pairing stops strengthening the conditioned response after roughly 1 minute of pairing, even if the shock intensity is below the behavioral saturation level. During conditioning, the odor is learned to predict the co-applied shock stimulus. As a consequence, the odor-evoked avoidance reaction stops strengthening at a level that depends on the shock strength, irrespective of the pairing time beyond 1 min. We found that associative synaptic plasticity, defined by a possibly nonlinear function of the CS-US correlation strength, as suggested by STDP models, fails to reproduce the early saturation of learning.

We suggest a simple phenomenological model for predictive plasticity according to which synapses change their strength proportionally to the prediction error. This error is expressed as a difference between the internal shock representation and the value representation of the odor. The model encompasses a description of the shock and value representation, the stochastic response behavior of individual flies, and the synaptic dynamics (using a total of 5 parameters). It faithfully reproduces our conditioning experiments (with a total of 28 data points from 3 different types of experiments) as well as previously studied trace conditioning experiments (without need for further fitting). As compared to the associative rules (Hebbian, linear and nonlinear STDP, covariance rule), the predictive plasticity rule obtained the best fits with the least number of parameters. We further compared the model by the Akaike information criterion that considers the number of parameters beside the fitting quality. This criterion yields a likelihood for the predictive plasticity rule to be the best one that is at least 7 orders of magnitude larger as compared to the other four associative rules we considered (see Table 1, Methods, and Figure 3-2).

**Table 1.** Akaike information criterion supports predictive plasticity

| Model $M$ | Predictive plasticity | Nonlinear STDP | Linear STDP | Covariance | Hebbian |
| --- | --- | --- | --- | --- | --- |
| Number of parameters | 5 | 10 | 8 | 6 | 5 |
| MSE($M$) | $6.40 \times 10^{-4}$ | $1.46 \times 10^{-3}$ | $1.45 \times 10^{-3}$ | $1.00 \times 10^{-2}$ | $1.24 \times 10^{-2}$ |
| -AIC($M$) | 114.45 | 81.36 | 85.55 | 35.48 | 31.46 |
| Relative likelihood $p$ | 1 | $6.49 \times 10^{-8}$ | $5.30 \times 10^{-7}$ | $7.15 \times 10^{-18}$ | $9.48 \times 10^{-19}$ |

### Error- versus target-driven predictive plasticity

The same phenomenological model of predictive learning may be implemented in two versions by the recurrent MB circuitry. In both versions the MBONs code for the odor value (‘valence’) that drives the conditioned response. For the error-driven predictive plasticity, the DANs directly represent the shock-prediction error by comparing the shock strength with its MBON estimate, and this prediction error modulates the KC-to-MBON plasticity (Figs 1B1 and 6A). For the target-driven predictive plasticity, the DANs represent the shock stimulus itself that is then provided as a target for the KC-to-MBON plasticity. In this target-driven predictive learning, the DANs may also learn to predict the shock stimulus based on the MBON feedback, preventing a fast extinction of the KC-to-MBON memory (Figs 1B2 and 6B).

Predictive plasticity for both types of implementation has its experimental support. In general, MBON activity is well recognized to encode the aversive or appetitive value of odors and to evoke the corresponding avoidance or approach behavior^4, 24, 30, 54, 55^, while KC-to-MBON synapses were mostly shown to undergo long-term depression, but also potentiation (see e.g.^50^). DAN responses are shown to be involved in both the representation of punishment and reward^6, 7, 26, 56^ that drive the aversive or appetitive olfactory conditioning^7^. This conditioning further involves the recurrent feedback from MBONs to DANs that may be negative or positive^33, 50^, see^5^ for a recent review. Moreover, the connectome from the larvae and adult fruit fly MBON circuit reveals feedback projections from DANs to the presynaptic side on the KC and the postsynaptic side on the MBONs at the KC-to-MBON synaptic connection^31, 32^, giving different handles to modulate synaptic plasticity.

With regard to the specific implementations, the error-driven predictive plasticity is consistent with the observation that DAN activity decreases during the conditioning^49, 51^. The two models have opposite predictions for learning while blocking MBON activity. The error-driven predictive plasticity would yield a higher LI, similarly as observed in^54^, while the target-driven predictive plasticity would yield a lower LI, similarly to^24^. It was also shown that some DANs increased their activity with learning while other DANs, in the same PPL1 cluster that is supposed to represent aversive valences, decreased their activity^51^. In fact, error- and target driven predictive plasticity may both act in concert to enrich and stabilize the representations. As shown in Figure 6, DAN activity would decrease in those DANs involved in error-driven and increase in those involved in target-driven predictive plasticity.

While error-driven predictive plasticity offers access to an explicit error representation in DANs, target-driven predictive plasticity has its own merits. If DANs and MBONs code for similar information, they can support a positive feedback-loop to represent a short-term memory beyond the presence of an odor or a shock, as it was observed for aversive valences in PPL1 DANs^6^ and for appetitive PAM DANs^33^. A positive feedback-loop between MBONs and DANs is further supported by the persistent firing between these cells after a rejected courtship that may consolidate memory of the rejection, linked to as specific pheromone^8, 57^.

### Distributed learning, memory life-time and novelty-familiarity coding

Target-driven plasticity has further functional advantages in terms of memory retention time. Any odor-related input to the DANs, arising either through a forward hierarchy from KC^48^ or a recurrence via MBONs to the DANs^6, 33^, will extend the memory life-time in a 2-stage prediction process: the unconditioned stimulus (*s*) that drives the DAN activity (*d*) to serve as a target for the value learning in the MBONs via KC-to-MBON synapses (*v* = *w*_MK_*o*), will itself be predicted in the DANs (see Eq. 11). Extending the memory life-time through circuit plasticity might be attractive under the light of energy efficiency, showing that long-term memory in a synapse involving de novo protein synthesis can be costly^8, 58^, while cheaper forms of individual synaptic memories likely have limited retention times. Moreover, distributed memory that includes the learning of an external target representation offers more flexibility, including the regulation of the speed of forgetting^45^.

Target-driven predictive plasticity may also explain the novelty-familiarity representation observed in the recurrent triple of KCs, DANs and MBONs^49^. The distributed representation of valences allows for expressing temporal components of the memories. Spontaneous activity in the KCs and their downstream cells^53^ injures a minimal strength of the KC-to-MBON synapses through predictive plasticity. A novel odor that drives KCs will then also drive MBONs and, to a smaller extent (as we assume), also DANs. If the DANs that represent the target for the KC-to-MBON plasticity are only weakly activated by the odor, the KC-to-MBON synapses learn to predict this weaker activity and depress. The depression results in a repetition suppression of MBONs and the corresponding familiarization of the fly to the ongoing odor. However, when the odor is cleared away, the MBON activity induced by spontaneously active KCs via depressed synapses now becomes lower than the spontaneous DAN activity, and predictive plasticity recovers the original synaptic strength. Eventually the spontaneous MBON and DAN activites match again (Eq. 13) and the response to the originally novel odor is also recovered, as seen in the experiment^49^.

Olfactory learning is likely distributed across several classes of synapses in the MB. The acquisition of olfactory memories was shown to be independent of transmitter release in KC-to-MBON synapses, although the behavioral recall of these memories required the intact transmission^59^. In fact, learning may also be supported by plasticity upstream of the MBONs such that the effect of blocking KC-to-MBON transmission during learning is behaviorally compensated. Predictive plasticity at the KC-to-MBON synapses requires the summed synaptic transmissions across all synapses in the form of the value *v* = *wo* to be compared with the target *d*, also during the memory acquisition. This type of plasticity would therefore be impaired by blocking the release.

### Distributed learning and absence of blocking

Distributed learning also offers flexibility in acquiring predictions from new cues. While the original Rescorla-Wagner rule would predict blocking^1^, this has not been observed in the fruit fly^46^. Blocking refers to the phenomenon that, if the first odor of a compound-CS is pre-conditioned, the second odor of the compound will not learned to become predictive for the shock. Because our predictive plasticity rules are expressed at the neuronal but not at the phenomenological level, predictions about blocking will depend on the neuronal odor representation. If the two odors activate the same MBONs, blocking would be observed since the MBONs are already driven to the correct value representation by the first odor. If they activate different MBONs, however, blocking would not be observed since the MBONs of the second odor did not yet have the chance to learn the correct value during the first conditioning. Hence, since blocking has not been observed in the fruit fly, we postulate that the odors of the compound-CS in these experiments were represented by different groups of MBONs.

### Concentration-specificity and relieve learning

How does our model relate to the concentration-specificity and the timing-specifity of odor conditioning? First, olfactory learning was found to be specific to the odor concentration, with different concentrations changing the subjective odor identity^60^. The response behavior was described to be non-monotonic in the odor intensity, with the strongest response for the specific concentration the flies were conditioned with. It was suggested that this may arise from a non-monotonic odor representation in the KC population as a function of odor intensity^35, 61^. Given such a presynaptic encoding of odor concentrations, the predictive olfactory learning in the KC-to-MBON connectivity would also inherit the concentration specificity from the odor representation in the KCs. Our predictive plasticity, and also the Rescorla-Wagner model, further predicts that learning with a higher odor concentration (but the same electroshock strength) only speeds up learning, but would not change the asymptotic performance.

Second, olfactory conditioning was also shown to depend on the timing of the shock application before or after the conditioning odor. While a shock application 30s after an odor assigns this odor an aversive valence, an appetitive valence is assigned if the shock application arises 30s before the odor presentation^16, 17, 62, 63^. Modeling the approaching behavior in the context of predictive plasticity would require duplicating our model to also represent appetitive valences, and the action selection would depend on the difference between aversive and appetitive valences. Inverting the timing of CS and US may explain ‘relief learning’ if a stopping electroshock would cause a decrease of the target for aversive MBONs (*d*^−^) and an increase of the target for appetitive MBONs (*d*^+^, see Eq. 12). An odor presented after the shock would then predict the increased appetitive target and explain the relieve from pain behavior, similarly to the model of relief learning in humans^64^.

Overall, our behavioral experiments and the plasticity model for the KC-to-MBON synapses support the notion of predictive learning in olfactory conditioning, with the DANs representing either the CS-US prediction error or the prediction itself. While predictive coding is recognized as a hierarchical organization principle in the mammalian cortex^65–68^ that explains animal^2^ and human behavior^69^ it may also offer a framework to investigate the logic of the MB and the multi-layer MBON readout network as studied by various experimental work^24, 32^.

## Methods and Materials

### Flies

We used *Drosophila* melanogaster of the Canton-S wild-type strain. Flies were reared on standard cornmeal food at 25°C and exposed to a 12:12 hour light-dark cycle. For the experiments groups of 60-100 flies (1-4 days old) were used.

### Behavioral experiments

The apparatus that was used to conduct the behavior experiment is based on^18^ and was modified to allow performing four experiments in parallel. Experiments were performed in a climate chamber at 23 – 25°C and 70-75% relative humidity. Training procedures were done in dim red light and tests were accomplished in darkness. Two artificial odors, benzaldehyde (Fluka, CAS No. 100-52-7) and limonene (Sigma-Aldrich, CAS No. 5989-27-5), were used for the experiments. 60*μl* of benzaldehyde was filled in plastic containers of 5mm and 85*μl* of limonene was filled in plastic containers of 7mm. Odor containers were attached to the end of the tubes. A vacuum pump was used for odor delivery at a flow rate of 7 *l*/min. Tubes lined with an electrifiable copper grid were used to apply electric shock. Shock pulses were 1.5 s long.

### Sequence shock experiments

Groups of flies were loaded in tubes lined with an electrifiable grid. After an initial phase of 90s, one of the odors was presented for 60s. At the same time electric shock pulses were delivered. After 30s of non-odorised airflow, the second odor was presented for 60s, without electric shock. Different electric shock treatments were used (see Fig. 2). In half of the cases benzaldehyde was paired with electric shock, while in the other half limonene was the paired odor. Whatever the idendity of the odor is, after pairing with the shock it is called the conditioned stimulus (CS+) while the odor paired with 0 shock strength is called the unconditioned stimulus (CS-). After the training flies were loaded into a sliding compartment and moved to a choice point in the middle of two tubes. Benzaldehyde was attached to one tube and limonene to the other. Flies could choose between the two odors for 120s. Then, the number of flies in each odor tube was counted.

### Repeated training experiment

One training block consists of 60s odor, 30s non-odorised air and 60s of the second odor. Four electric shock pulses were delivered after 15, 30, 45 and 60s of the first odor presentation. Flies were exposed to this training block one, two or four times. The time between the training blocks was 90s. For ‘0.5 repetitions’ (as reported in Fig. 4) only two pulses were delivered 45 and 60s after onset of the odor and this block was not repeated. Experiments were performed with electric shock pulses of 25 and 50V. After the training, learning performance was tested as in the sequence shock experiment.

### Continuous shock experiments

Continuous electric shock was used to train the animals instead of pulses. Electric shock was applied during the entire presentation of the first odor (odor X). odor X and shock duration were 10, 15, 30, 45, 90 or 120s. The second odor (odor Y) was presented for the same duration as odor X and the electric shock. odor Y was always applied 30s after the end of odor X presentation. Experiments were performed with 25 and 50V. The learning test after the training was identical to the sequence shock experiment.

### Minimal shock detection

For the electric shock avoidance tests, flies were loaded into a sliding test chamber (compartment). The chamber with the flies was pushed to a choice point between two arms (tubes) with an electrifiable grid at the floor. The grid in one tube was connected to a voltage source (of strength *S*), whereas the other was not. Electric shock was delivered continuously for 30s and then the number of flies in each tube was counted. For a shock of strength *S* = 5, 9 and 12.5V we measure a performance index PI(*S* = 5V) = 0.006 ± 0.014 (mean ± standard error of the mean, SEM), PI(*S* = 9V) = 0.030 ± 0.014 and PI(*S* = 12.5V) = 0.068 ± 0.019, respectively. For *S* = 7V we estimated the mean PI to be roughly 0.01, with a SEM to be roughly twice as large, 0.02, see Figure 2-1.

### Parameter optimization

The parameters are optimized to minimize the least square error between the experimental data and the model simulation. The optimization is done in Matlab (R2014a), using Interior point method with maximum 3000 iterations, 1.0e-06 tolerance. Initial conditions of the parameters are uniformly sampled from a wide interval, and all optimized parameters with similar overall performances were clustered around the ones reported in the caption of Fig. 3. The same set of parameters for the predictive plasticity (Eq. 7) is used throughout. The *mean square error* (MSE) between data mean and model mean is calculated by summing the squared error of the means (with the same *N*_fly_ and *N*_trial_) for all 28 data points across all experiments, divided by 28. The parameters for the predictive learning model are reported in the caption of Fig. 3, the ones for the other models below.

### Adaptable learning rate

The learning rate is assumed to increase with increasing stimulus strength 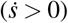 and otherwise passively decays. Its dynamics has the form

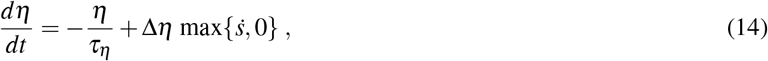

with optimized parameters *τ_η_* and Δ*η*. We were choosing *τ_η_* = 133.48s and Δ*η* = 0.057 in all the experiments using predictive learning rule except for the simulation of the target-driven learning model in Figure 6B where we set *τ_η_* = 26.7s and Δ*η* = 0.74. For the discrete time simulations, a step-increase Δ*s* in the shock stimulus triggers a step increase in *η* by Δ*η* Δ*s*.

### Linear STDP, nonlinear STDP, and covariance rule

Given the analogy of the olfactory conditioning to spike-timing (or stimulus-timing) dependent plasticity (STDP)^7, 13, 16, 17^, we considered two different forms of STDP rules. The linear STDP learning rule is

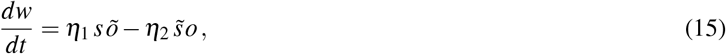

where 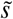 is the low-pass filtered *s* with filtering time constant *τ_s_* = 17.87*s*, *õ* is the low-pass filtered *o* with time constant *τ_o_* = 7.47*s*, *η*_1_ = −0.47, *η*_2_ = −0.47, *α* = 0.23, and *S*_∘_ = 9.31*V*. For the linear STDP we get MSE = 5.489 × 10^−3^ for the indicated optimized parameters.

The nonlinear STDP rule is of the form

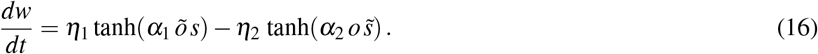

The learning rates *η*_1/2_ were allowed to be both positive and negative. The optimal parameters are *η*_1_ = 0.01, *η*_2_ = 0.19, *τ_o_* = 51.20*s*, *τ_s_* = 124.12*s*, *α* = 9.93, *α*_1_ = 9.93, *α*_2_ = 0.44, *S*_∘_ = 11.91*V*, with a MSE = 6.550 × 10^−3^.

The covariance rule has the form

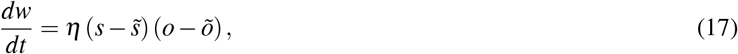

and the optimal parameters are *η* = 0.12, *τ_o_* = 300.00*s*, *τ_s_* = 19.18*s*, *α* = 0.53, *S*_∘_ = 9.13*V*, with a MSE = 8.236 × 10^−3^.

All these 3 rules (Eq. 15–17) failed mainly in reproducing the repetitive conditioning experiments (Fig. 4B), see Supplementary Material. Overall, the MSE for all these associative rules is roughly 10 times bigger than the MSE for the predictive rule (MSE = 6.393 × 10^−4^).

### Associative learning rules with adaptive learning rate

We also tested the associative learning rules with the adaptive learning rate from Equation 14. Although the MSEs get smaller for both linear and nonlinear STDP rules, they remain twice as large as for the predictive learning rule (see Figure 3-2). The covariance rule (with optimised parameters *α* = 0.12, *S*_∘_ = 3.68, *τ_o_* = 37.70*s*, *τ_s_* = 1498.38*s*, *τ_η_* = 60.54*s*, Δ*η* = 1.00, with a MSE = 1.00 × 10^−2^) did not profit from the adaptable learning rate. For the linear STDP learning rule (Equation 15) the optimal parameters with adaptable learning rate are *α* = 0.31, *S*_∘_ = 3.20, *τ_o_* = 8.29*s*, *τ_s_* = 171.05*s*, *τ_η_*_1_ = 16.01*s*, *τ_η_*_2_ = 5.78*s*, Δ*η*_1_ = 0.39, Δ*η*_2_ = 5.71, with a MSE = 1.45 × 10^−3^.

For the nonlinear STDP learning rule (Equation 16), the optimal parameters are *α* = 0.13, *S*_∘_ = 3.21*V*, *τ_o_* = 8.30*s*, *τ_s_* = 171.10*s*, *τ_η_*_1_ = 16.02*s*, *τ_η_*_2_ = 5.82, Δ*η*_1_ = 8.96, Δ*η*_2_ = 5.14, *α*_1_ = 0.24, *α*_2_ = 6.05, with a MSE = 1.46 × 10^−3^. For the Hebbian additive rule in Equation 6 with adaptable learning rate, the optimal parameters are *α* = 0.05, *S*_∘_ = 5.08, *τ_o_* = 1.50*s*, *τ_η_* = 49.81*s*, Δ*η* = 5.46, with a MSE = 1.24 × 10^−2^.

### Model comparison based on the Akaike information criterion

We compared the various models on the basis of the Akaike information criterion (AIC) that puts the model accuracy on the data set into relation to the number of parameters used to achieve this accuracy^70, 71^. Assuming that the estimation errors of all *n* experimental conditions are normally distributed with zero mean, the AIC for a given model *M* is calculated as a log-likelihood,

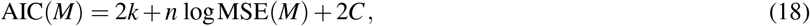

where *k* is the number of parameters in the model, *n* = 28 the number of experimental conditions, and 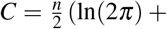 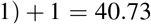. The relative likelihood *p* for model *M* to be true as compared to the predictive plasticity model *M*_0_ is 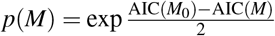 (Table 1).

## Data availability

The mathematical model (Matlab) including the experimental data will be available on https://github.com/unibe-cns.

## Acknowledgments

We thank members of the Senn and Sprecher lab for helpful support, and in particular Robert Urbanczik^†^, Martin Wiechert for fruitful discussions and Elena Kreutzer, Dominik Spicher, Jakob Jordan, Rui Ponte Costa, Ashok Litwin-Kumar and Emmanuel Perisse for helpful comments on the manuscript. WS is grateful for inspiring conversations with an anonymous reviewer. This research was supported by the SystemsX.ch initiative (for SS and WS, grant 51RT-0-145733, evaluated by the SNSF), the SNSF (grant 310030L-156863, WS), a SNF sinergia grant (CRSII5-180316 led by F. Helmchen) and a grant from the China Scholarship Council (201408080117, CZ). WS and MAP were supported by the European Union’s Horizon 2020 Framework Programme for Research and Innovation under the Specific Grant Agreement No. 720270, 785907, 945539 (Human Brain Project SGA 1-3, related to predictive coding).

## Author contributions statement

WS, SGS, CZ and YFW designed the experiments. WS and CZ developed the mathematical model, YW and SD performed the experiments, CZ and MAP performed the computer simulations, and CZ, WS, YW and SGS wrote the manuscript.

## Supplementary Information

**Figure 2-1.**
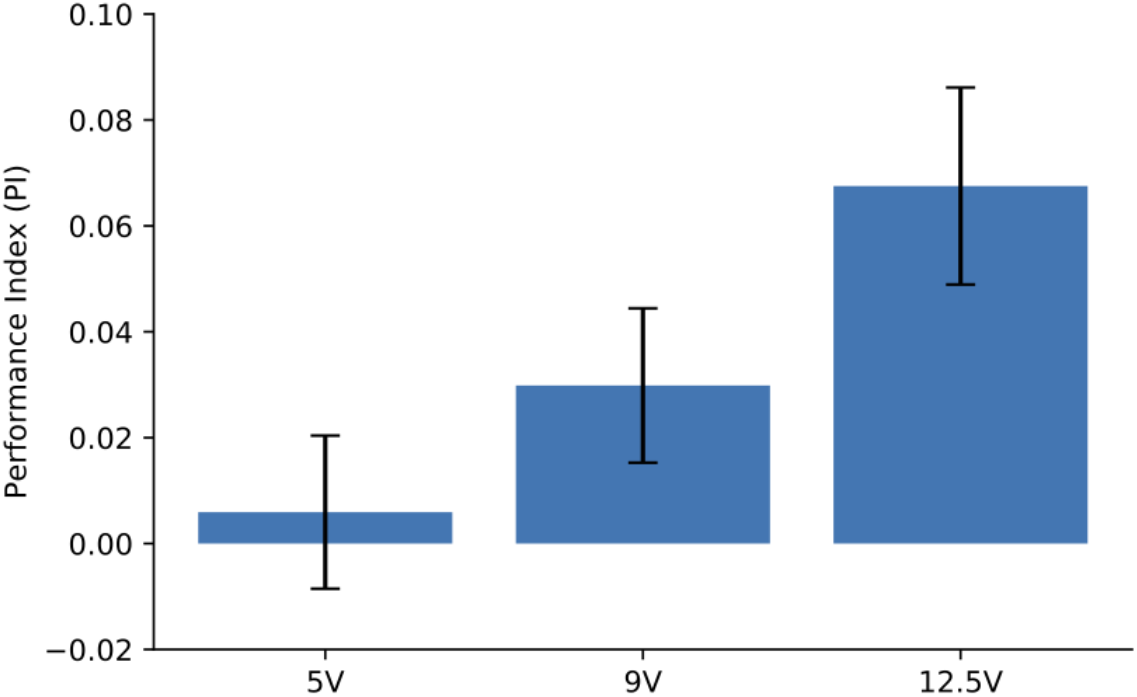
The minimal detectable shock stimulus: Electrical shocks with 5V, 9V, and 12.5V are applied in the electric shock avoidance tests. The performance index decreases as the shock intensity decreases. PI is close to 0 when the shock intensity is 5V and significantly above 0 for 9V, and we estimate *S*_∘_ ≈ 7V. The error bars represent the SEM.

**Figure 3-1.**
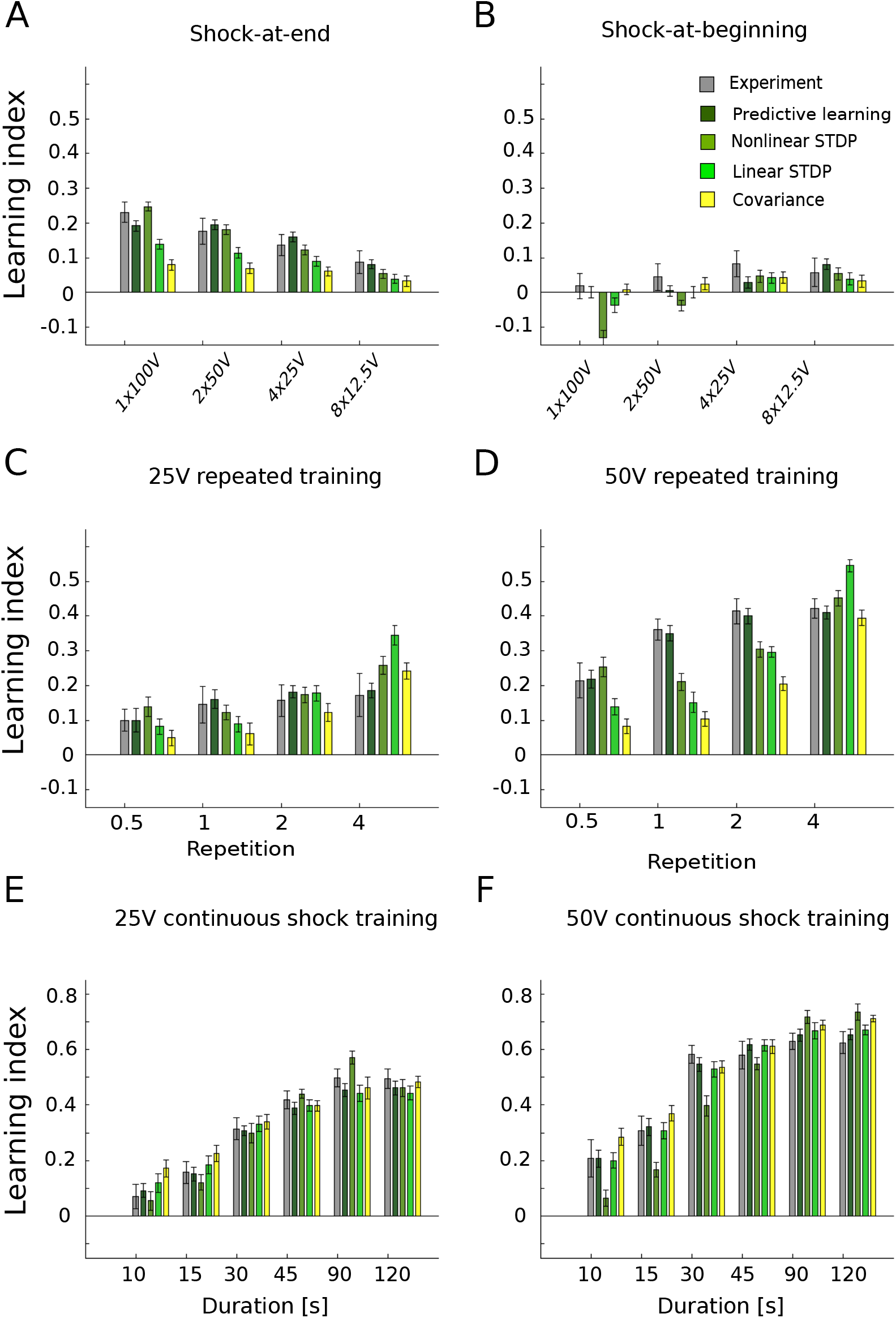
Associative learning rules are not able to fit all experimental data. A comparison of all learning rules. **(A)** Temporal sequence training with shocks-at-end alignment. **(B)** Temporal sequence training with shocks-at-beginning alignment. **(C)** Repeated training with 25V. **(D)** Repeated training with 50V. **(E)** Continuous shock training with 25V. **(F)** Continuous shock training with 50V. The associative (linear and nonlinear STDP and covariance) learning rules fail mostly in the repeated training experiments, as they are not able to capture the saturation in the experimental data.

**Figure 3-2.**
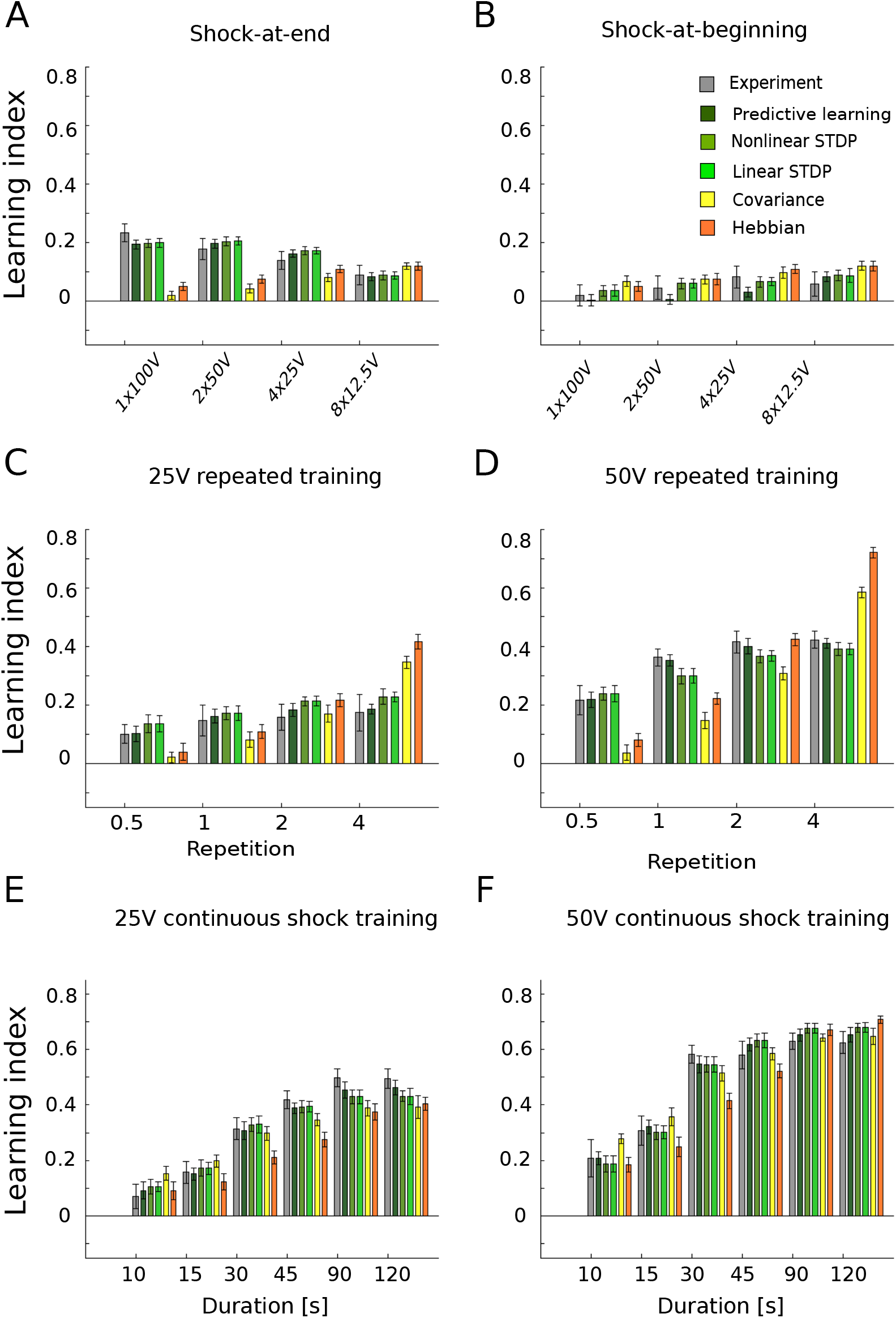
Associative learning rules with adaptive learning rate. A comparison of all learning rules. **(A)** Temporal sequence training with shocks-at-end alignment. **(B)** Temporal sequence training with shocks-at-beginning alignment. **(C)** Repeated training with 25V. **(D)** Repeated training with 50V. **(E)** Continuous shock training with 25V. **(F)** Continuous shock training with 50V. The covariance rule and simple Hebbian rule is not able to reproduce all the data. The linear and nonlinear STDP rules perform better with adaptive learning rate, but still have twice as big as MSE comparing to the predictive plasticity rule

**Figure 4-1.**
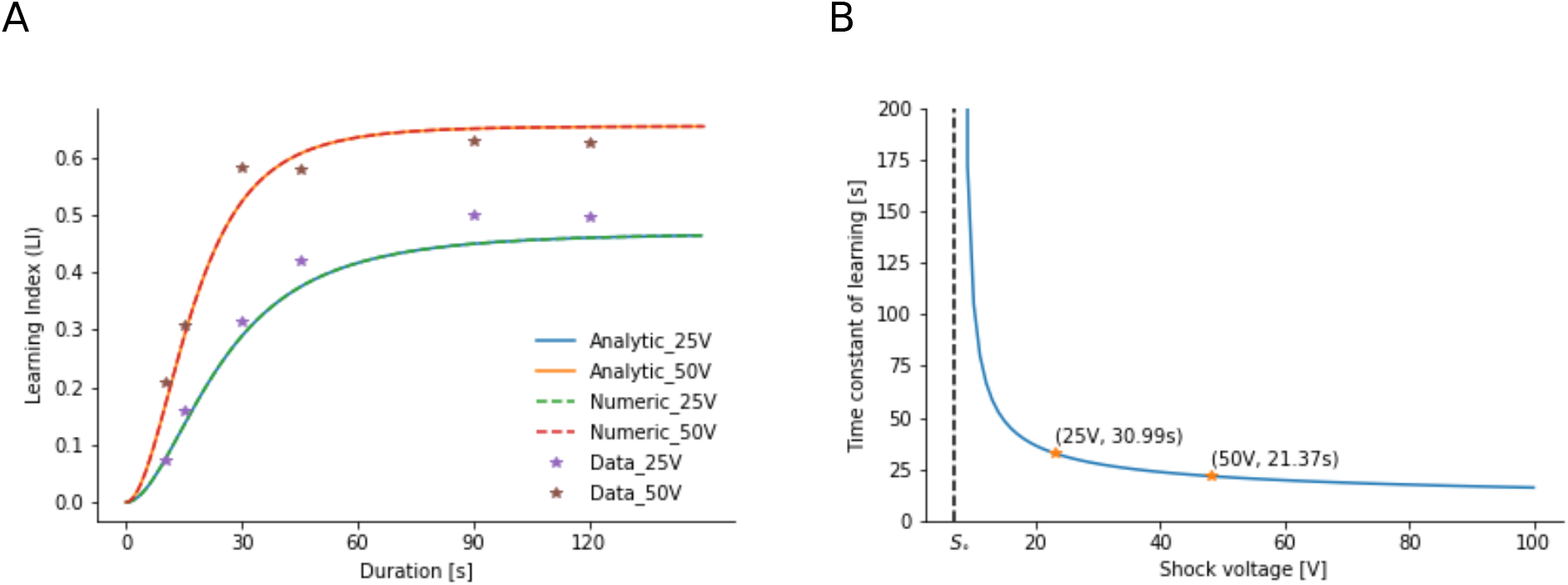
Extracting the learning time constant for the ongoing shock experiments. **(A)** The analytical solution (solid lines, Equation S5) for the development of the LI matches the numeric simulation (dash lines, overlaid) for the ongoing conditioning experiments (stars). **(B)** The time constant of learning diverges for shock intensity *S* close to *S*_∘_, and it monotonically decreases for shock intensities beyond *S*_∘_. For 25V, the learning time constant is 30.99s; for 50V, it is 21.37s (with optimized parameters from the model, see caption of Fig. 3).

## Analytical solution for the ongoing shock experiments

In the ongoing shock experiments, the odor and shock stimuli are both turned on for the whole pairing duration, and turned off when pairing stops. For a constant odor concentration *o* = 1 for *t* ≥ 0 while *o* = 0 before, the dynamics of the odor eligibility trace, 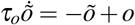, is solved by

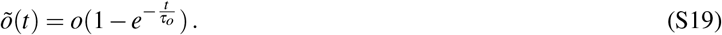

With a step increase of the shock from 0 to Δ*s* at time *t* = 0, the learning rate *η* according to the dynamics Equation 12 undergoes a step increase by Δ*η*Δ*s* that again decays during the constant voltage application,

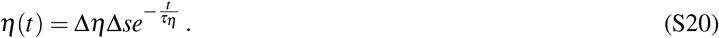

The weight *w* from the KCs to the MBONs Further, according to the predictive plasticity rule Equation 7, 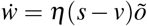, exponentially increases from 0 to 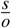,

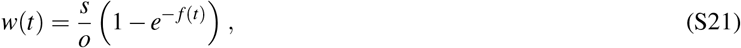

with

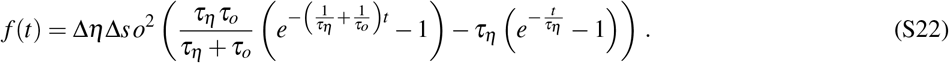

Plugging Equation 19 into *v* = *wo* and this into expression for the avoidance probability *p_cs_*(*v*), Equation 5, the learning index LI(*v*) = 2*p_cs_*(*v*) − 1 develops in time according to

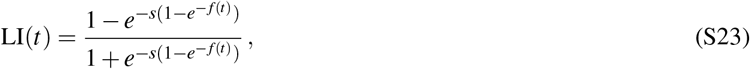

and this converges to LI(*s*) is as in Equation 8.

